# Environmental cues influence timing and location of construction activity in a beaver damming complex

**DOI:** 10.1101/2024.10.21.619304

**Authors:** Jordan Kennedy, Helen McCreery, Audrey Jones, Dannette Spotted Horse, Candice Chen, Emily Fairfax, Justin Werfel

**Author notes:** JK: Outer Coast College, Sitka, AK 99835, USA HM: Tufts University, Medford, MA 02155, USA AJ: FlowWest, Oakland, CA 94604, USA DSH: New Day Ranch, Billing, MT 59101, USA CC: Massachusetts Institute of Technology, Cambridge, MA 02139, USA EF: University of Minnesota, Minneapolis, MN 55410, USA JW: Harvard University, Cambridge, MA 02138, USA.

## Abstract

Beavers are famous for building extensive damming complexes^1^, which can extend over kilometer-scale distances^2^ and persist for centuries^3^. Dams have major impacts on their surrounding environment, including effects on the hydrology, geomorphology, and ecosystems of the area^4^. Current understanding of how the environment in turn affects beavers’ building activity centers on the single auditory cue of running water, based primarily on captive studies^14,15^. Observational challenges have limited a more detailed picture of the full feedback loop between beavers and their environment. Here we describe the detailed progress of new construction by 20 beaver colonies in northwest Montana, through field studies using drones to obtain surveys with spatial and temporal resolution each three orders of magnitude finer than typically reported^5-13^. We show that both the timing and location of beaver construction activity are influenced by environmental factors. Initiation of trail clearing, a stage preceding dam building, was associated with a narrow range of stream flow rates. Beavers preferentially built in locations associated with preexisting canals. These results emphasize the importance of non-dam elements in the beavers’ construction and its coordination across the colony, and point to environmental feedback processes that may span across years and unrelated colonies.

---

To cope with substandard hydrologic conditions that do not meet their life-cycle requirements, North American beavers (*Castor canadensis*) construct extensive damming complexes (Fig. 1). These complexes emerge over considerable spatial (100s to 1000s of meters) and temporal (decades to centuries) scales, and fulfill a variety of functional roles for the colony. Elements of a beaver complex (Fig. 1A) include the iconic dams (often several per colony), ponds that flood upstream of dams, a lodge or bank den where the colony lives, trails cleared over land as beavers forage terrestrially, canals that beavers excavate to help access and transport vegetation via aquatic pathways, food caches where woody vegetation is submerged in the ponds for later consumption, and earthen scent mounds marking territorial boundaries.

**Figure 1.**
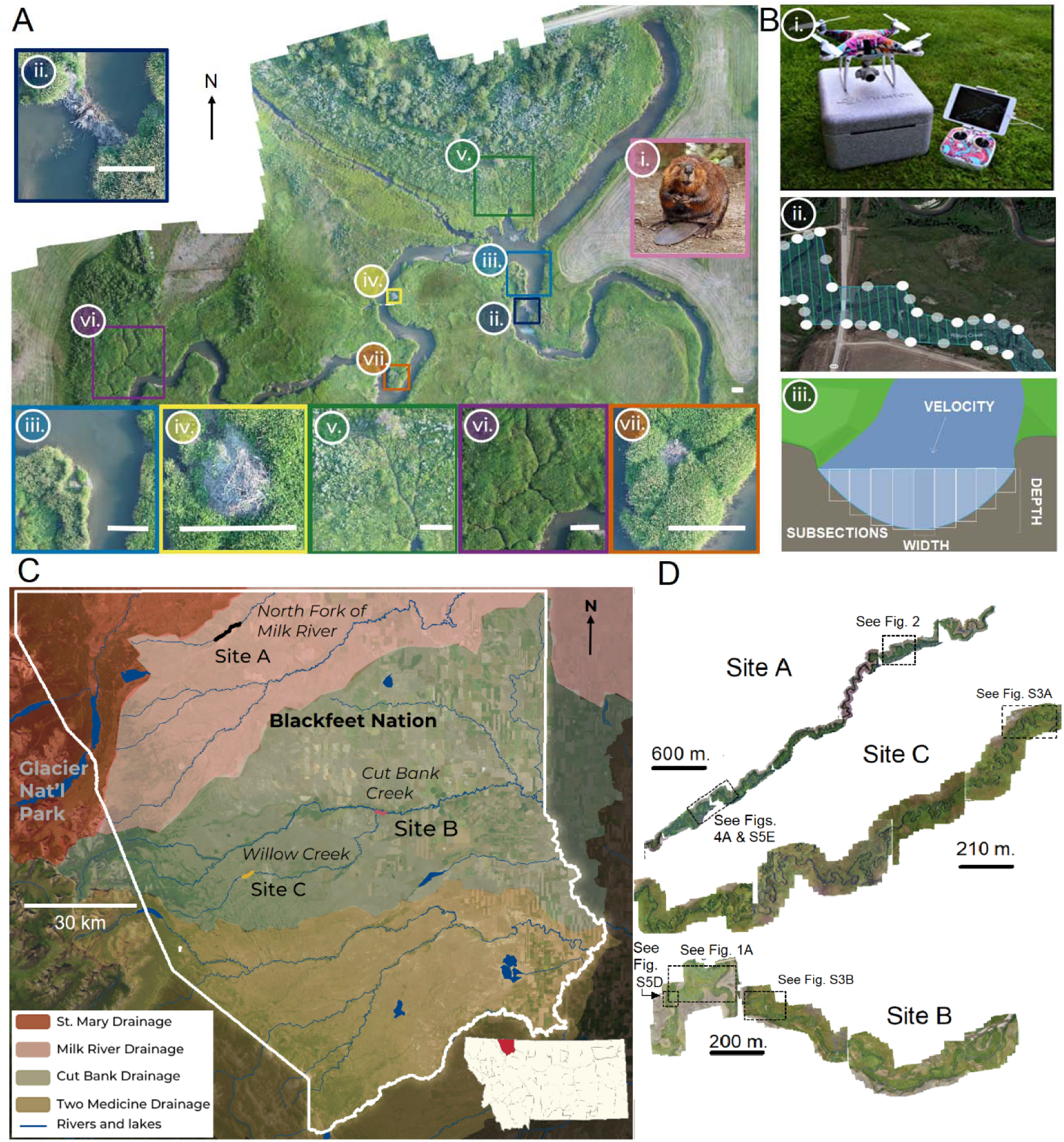
Study area and methods overview. (**A**) Orthomosaic of a damming complex built by one beaver colony (designated B1 in Fig. 3 and the SI), created from photos taken via UAV survey. Features: (i) beaver; (ii) dam; (iii) pond; (iv) lodge; (v) trails; (vi) canals; (vii) scent mound. Scale bars = 5 m. (**B**) The imagery was captured using (i) a DJI Phantom 4 flown in (ii) a Nadir flight plan. (iii) Stream discharge measurements were taken upstream and downstream of the field sites using the area-velocity method^15^. (**C, D**) The study included three field sites on two watersheds, with a total of 20 identified beaver colonies.

Studies of beaver construction have typically focused on the impacts of dams and ponds on the surrounding environment^4^. Less studied are the impacts of the other built elements of the complex, and the impacts of environmental factors on beavers’ decisions of when and where to build. These lacunae are due in part to challenges of observation: large-scale observations of habitat modification are typically performed with aerial photography at low spatial (∼30 m) and temporal (10+ years) resolution^5-13^; beavers are nocturnal and semi-aquatic, and range over territories of several kilometers, making their activity difficult to observe directly; and initiation of new dams can be a rare event, hard to capture by chance. Consequently, large-scale observations may exclude smaller features such as trails and canals, as well as details of the dam construction process, and knowledge of construction behavior is largely based on observations of captive beavers. These captive studies have highlighted the sound of running water as a cue the animals use for repairing and maintaining dams^14,15^. Cues beyond that auditory one, and how they influence when and where beavers choose to engage in construction in a natural setting, have not been explored.

We used unmanned aerial vehicle (UAV) photogrammetry to conduct observations of 15.9 km of three rivers in northwest Montana, encompassing the territories of 20 beaver colonies. We captured images at high resolution in both space (∼1.4 cm/pixel) and time (∼weekly), and measurements of river flow, over four months in the summer of 2018. In this region, annual snowmelt typically washes out most of the previous year’s dams, forcing beavers to recapitulate the dam-building process each year. The study’s goals were to observe how beavers’ construction activity is influenced by hydrologic conditions, landscape features, and other environmental factors. Our hypotheses were that the timing of initiation of dam construction would be driven by stream flow falling below a threshold, and that the location of novel dam building would be driven by natural landscape features such as the sinuosity and width of the river.

## Monitoring beaver dam construction

We selected three field sites within the Blackfeet Indian Reservation (BIR) boundary (Fig. 1C, D), each a continuous length of incised streambed with beaver-occupied lodges and a known history of dam building. In this area, the low-lying woody vegetation allows for a relatively unobstructed view of the waterways while providing enough material for beavers to build and sustain colony populations, making the sites particularly amenable to monitoring by aerial photography.

We surveyed each site approximately once per week from May 2018 to August 2018 (Supplementary Table 2), using DJI Phantom UAVs executing predefined flight paths (Fig. 1B) for consistency across observations. The UAVs collected high-resolution RGB photographs (1.3– 1.7 cm/pixel) which were used to create 2D orthomosaics and 2.5D surface models of each site at each time. The orthomosaics were aligned with ground-truth maps using Geographic Information Systems (GIS) software. Damming complex features (dams, lodges, trails, canals, scent mounds, and riverbanks) were manually identified and annotated for each orthomosaic. We identified 20 colonies in total across the three field sites by their lodges, and associated the damming complex features with colonies based on proximity and known information about typical territory sizes^17^. A single additional UAV scan was taken at all sites in September 2019.

Riparian vegetation is critical for beaver colonies for both building materials and food. As a measure of vegetation quality throughout the season, we used the RGB images to calculate the Green-Red Vegetation Index (GRVI) within the riparian corridor, which serves as a proxy for photosynthetic productivity^18^, for each site and time point. Other RGB-derived greenness phenologic indices are compared in Extended Data Fig. 7.

In addition to visual records of the sites, we measured stream flow at the upstream and downstream end of each site throughout the season, using the velocity-area method^16^ (Fig. 1Biii).

## Trail initiation associated with flow

The construction of a new damming complex results in significant aquatic and terrestrial alterations (Fig. 2, Extended Data Fig. 5). Aquatic alterations, extensively documented in previous studies^4^, are a direct consequence of dam construction, pond formation, and canal development. Terrestrial modifications are the result of foraging and trail-clearing activities. The intricate network of trails and canals provides beavers with access to terrestrial and aquatic resources. Although long-term environmental modifications resulting from beaver activity have been well documented based on observations 10 or more years apart^11,19^, it is notable that this study reveals substantial changes that occurred in just 12 weeks.

**Figure 2.**
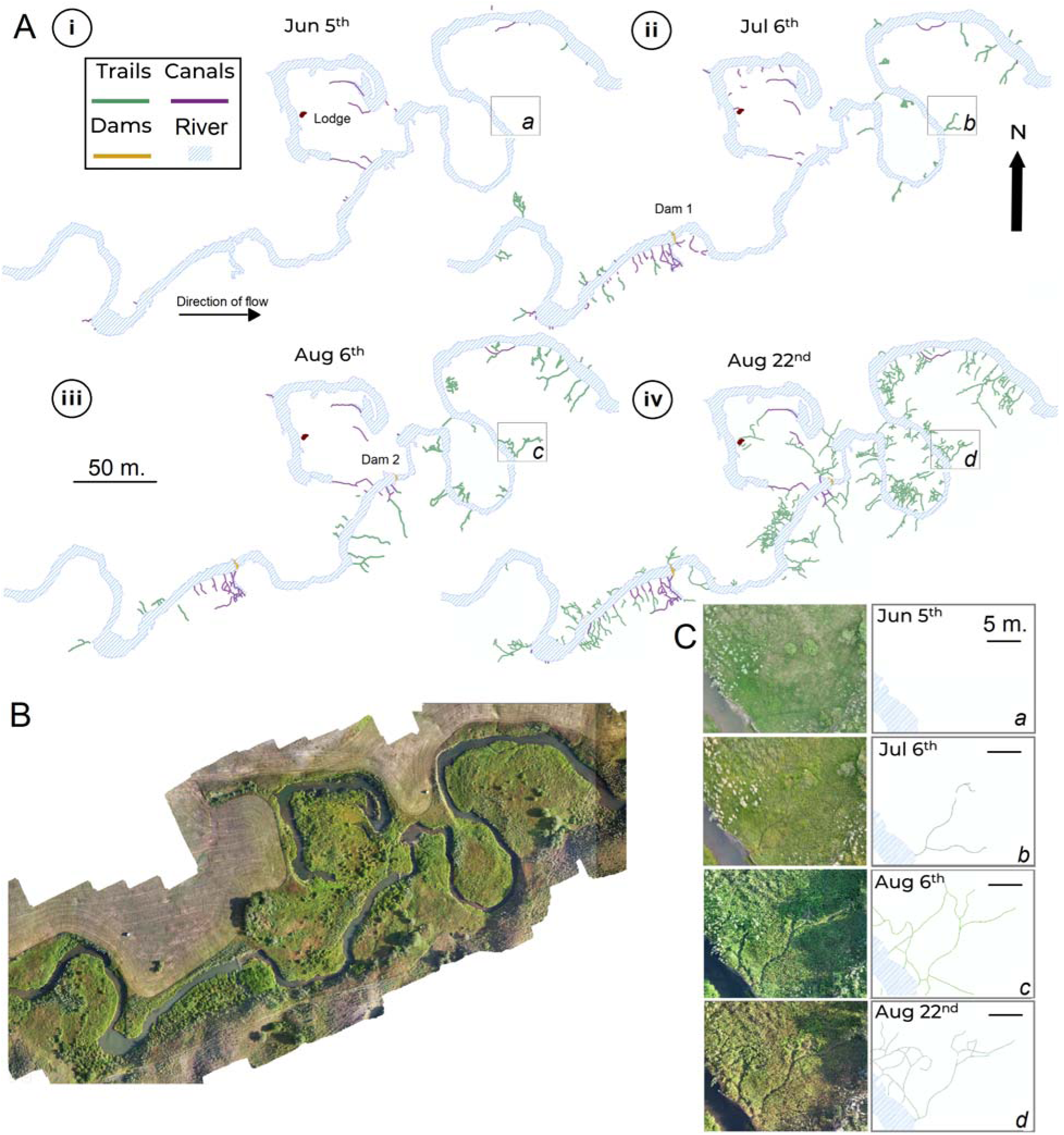
Seasonal development of a damming complex. (**A**) State of the territory of Colony A1 at four time points (i–iv), demonstrating dam, canal, and trail growth. Lettered boxes show the locations of the closeups in panel C. (**B**) UAV-based orthomosaic image of the territory. (**C**) A close-up of a developing trail system at four time points.

Of the 20 identified colonies, nine had at least one complete dam that survived after the spring snow melt, while 11 had all dams completely or partially washed out and thus needed to engage in dam construction. Therefore, we focused on the latter colonies in investigating environmental influences on the timing of new construction. Of these 11 colonies, we further excluded from this analysis two that abandoned their sites early in the season and one that arrived late in the season, to consider only those eight colonies present and building during the full period of observation (Supplementary Table 3).

For each damming complex, we identified the date on which construction was first observed, for both trails and dams. Across the sites and colonies, the dates of construction initiation spanned several weeks (Fig. 3A). Trail construction consistently preceded dam building. While (contrary to our original hypothesis) there was no consistent relationship between dam initiation and flow rate, trail initiation was associated with a narrow range of flow for all colonies (Fig. 3C).

**Figure 3.**
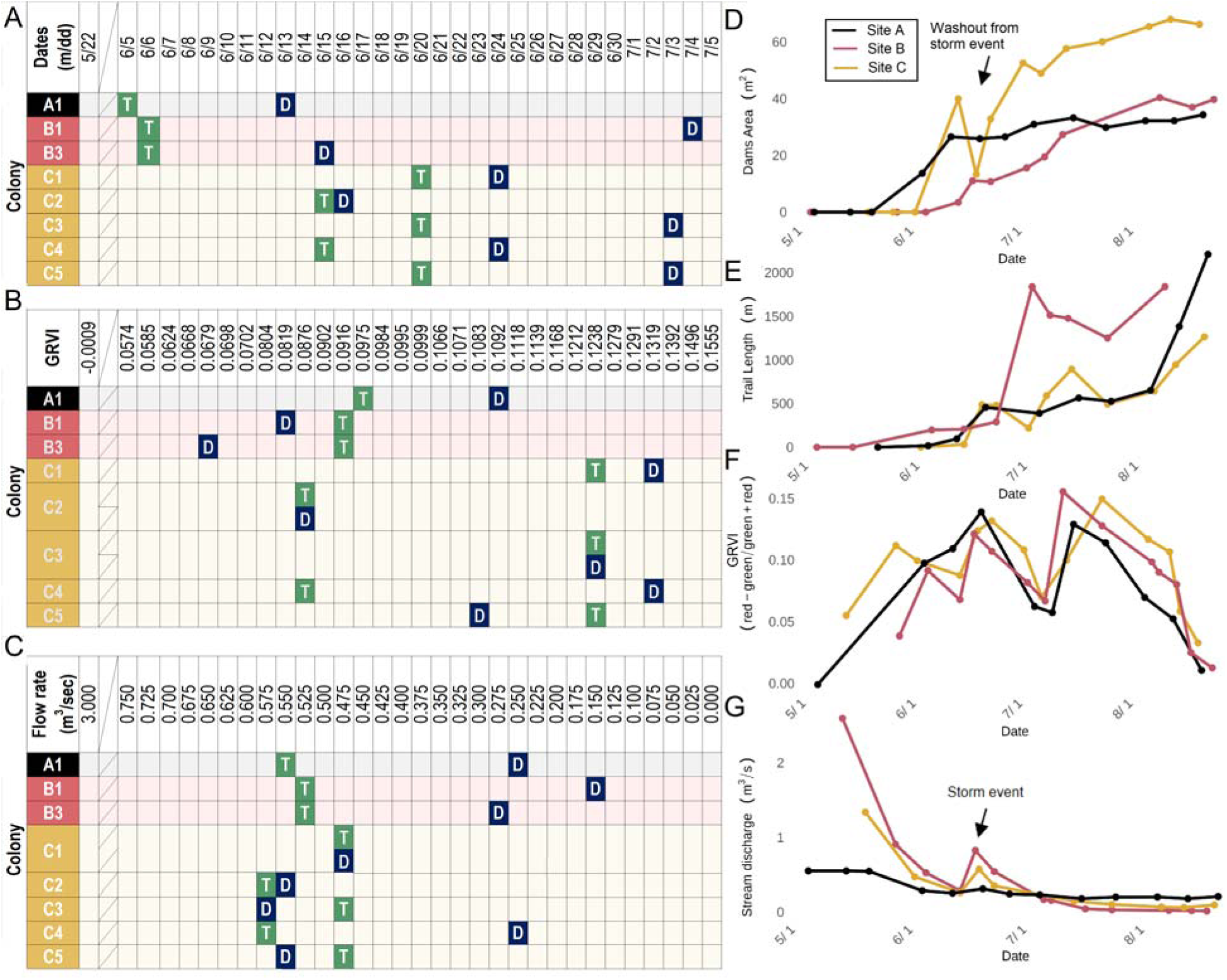
Dam and trail construction and environmental factors. (**A**) Dates on which construction of trails (T) and dams (D) was first observed for each colony. (**B**) Vegetation index (GRVI) value when construction was first observed. (**C**) Stream flow rate when construction was first observed. (**D**) Total area of all dams for all colonies at each field site over time. The sharp decrease for site C in late June corresponded to a storm surge event (see panel G) that washed out some dam construction. (**E**) Total length of all trails for all colonies at each field site over time. (**F**) Average GRVI value for each field site over time. (**G**) Average stream volumetric flow rate for each field site over time.

We used exact multinomial tests (EMT) of goodness-of-fit to assess statistical strength of the associations with date, vegetation index, and flow rate for both trail and dam initiation (see Materials and Methods). The proportions of both dams and trails initiated across values of the vegetation index did not significantly differ from the null expectation (for dams: EMT, 45 events, *p* > 0.18; for trails: EMT, 45 events, *p* > 0.33). The association between flow rate and construction initiation was significant for trails (EMT, 1287 events, *p* < 0.00028) but not dams (EMT, 1287 events, *p* > 0.19). Proportions of initiations across dates significantly differed from the null expectation for trails (EMT, 495 events, *p* < 0.0060), but not for dams (EMT, 495 events, *p* > 0.11). Despite the challenge of disentangling flow rate from date, it is important to note that where these metrics differed, beaver colonies initiated trail building at a consistent *flow rate* and *not* at a consistent date. At every colony, beavers began to construct trails coincident with a local flow rate in the range of 0.475–0.600 m^3^/s, even though the associated dates spanned several weeks. We conclude from this that flow rate, not date, is the primary factor cuing the start of trail building.

## Canal proximity influences trail location

We next investigated factors influencing where beavers choose to build. Our initial focus was on new dams, and geographical features of the stream that might make a site attractive, in particular its width and sinuosity.

In all, we observed 46 dams constructed in 2018 by the 20 colonies we surveyed (Extended Data Fig. 8A). Of these, 26 were cases of rebuilding a dam from the previous year that had been breached by the snowmelt but for which some trace remained; 19 were newly constructed from scratch, where no dam had been the previous year; and one was built on a natural logjam. Additionally, 7 dams from the previous year that were breached by snowmelt were not rebuilt. We also identified 31 “historic” dams that were present the previous year and left intact after the snowmelt; based on Google Earth imagery from previous years, such dams could persist for years or decades.

For the 19 dams built from scratch, we used a resampling analysis (see Materials and Methods) to generate a null distribution of mean stream widths that would be expected if new dams were constructed at random locations within the colonies’ territories (Fig. 4Bi). We performed the same procedure for sinuosity (Fig. 4Bii). These analyses found that the actual mean values fell well within these distributions. Thus our observations showed no evidence that those geographical features have any effect on locations selected to build dams. Inspection of Google Earth imagery across three decades^20^ revealed that canal architectures develop on a time scale of multiple beaver generations (Fig. 4A), while trails do not persist from one year to the next. As canals facilitate the transport of woody vegetation^21^, another possible factor in the locations chosen for new dam locations is proximity to previously existing canals.

**Figure 4.**
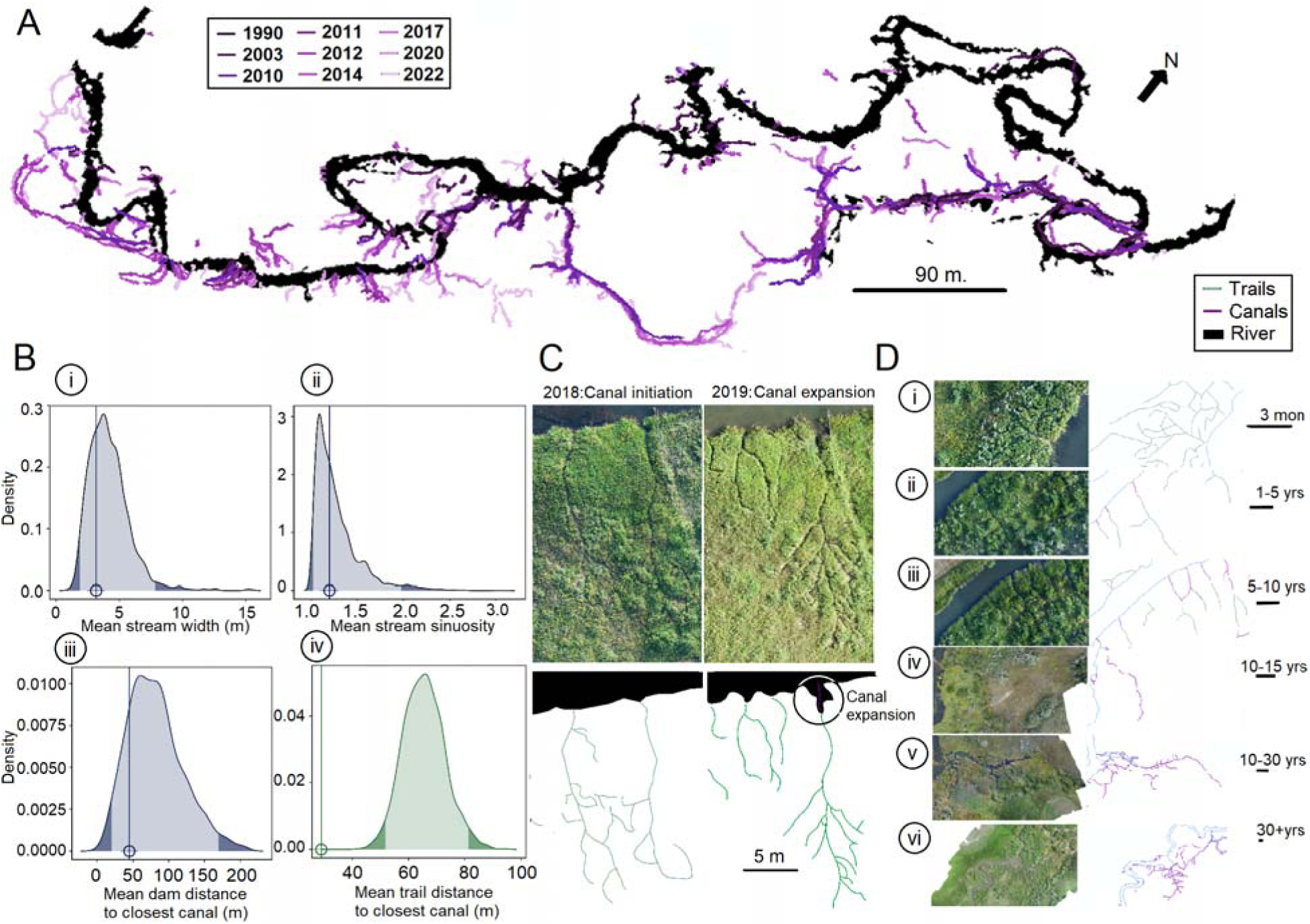
Dam and trail location analysis and longer-term canal network dynamics. (**A**) Manually annotated canals from Google Earth imagery from 1990 to 2022^20^. Darker purple corresponds to older canals, lighter purple to newer canal excavation. Canals increase in length and complexity over the 32-year observational window, ramifying and sometimes eventually interconnecting with others nearby; similarly, canals can shrink as branches are abandoned and silt up. (**Bi - iv**) Null distributions for different quantities based on resampling analyses, each based on 1000 simulated experiments in which the location of structures is randomized (see Materials and Methods). Dark gray shading indicates the most extreme 2.5% at each tail of each distribution. Actual observed values are marked as circles with vertical lines; for (i)–(iii), the observed value falls well within the null distribution, while for (iv), the observed value is far into the tail. (i) Mean stream width at dam locations. (ii) Mean stream sinuosity at dam locations. (iii) Mean distance from dams to the nearest canal. (iv) Mean distance from trails to the nearest canal. (**C**) UAV scans from 2018 and 2019 show the expansion of a canal initiated at the root of a large and active trail network. (**D**) Hypothesized evolution of a trail to a canal system, with speculative time scales for each stage at right. All images are taken from site A. (i) Trails are cleared. (ii) A heavily utilized trail system begins to become a short canal. (iii) Canals are extended. (iv) Canals replace a trail almost entirely. (v) Canals become complex with branches forming. (vi) Canals begin to interconnect with other canals, creating a complex aquatic network. Scale bars: 5 m.

To test this second hypothesis, we measured the distance along the river from each newly constructed dam to the nearest preexisting canal and compared the mean distance to a null distribution using a resampling analysis (see Methods). The result did not support the hypothesis that canals influence new dam locations (Fig. 4Biii). Because the number and density of canals differ among the colonies’ territories, the expected average distance from dams to canals also varies among territories. To ensure that the result from the pooled analysis was not an artifact of variation among colonies, we therefore also performed this analysis colony-by-colony, obtaining similar results (Extended Data Fig. 10A). Because of the salience of trail construction suggested by the timing analysis, we performed the same analysis for proximity of trails to canals. This analysis, by contrast, showed that trails are built much closer on average to preexisting canals than would be expected for random placement, both when all trails constructed by all 20 colonies were considered together (Fig. 4Biv; *p* < 10^−8^) and when the analysis was performed on a colony-by-colony basis (Extended Data Fig. 10C).

For 19 of the 20 colonies we observed, the colony was already present in its territory at the start of the season. Building in the immediate vicinity of canals thus could correspond to repeating construction in an area previously successful for the colony. However, it is important to note that the utility of canals is to facilitate access to terrestrial and aquatic resources^21^. Canals allow beavers to easily transport woody vegetation required for building dams, lodges, and food caches. An area of vegetation exploited by beavers one year may remain denuded the following year, regrowing only at a time scale of multiple years (Extended Data Fig. 5D,E). Therefore building near a canal is not necessarily synonymous with building near resources. Rather, canals may serve as an indication for beavers that a location has been productive at *some* point in the past and is worth investigating as a potential construction site.

Consistent with this interpretation, the one colony we observed that migrated into the study area mid-season from elsewhere (colony B2; Extended Data Fig. 3B, 6A,B) settled in a region with preexisting canals left by a previous colony. Moreover, although colonies frequently relocate^22^, the presence of beavers is closely associated with that of canals: we observed 387 canals in 6334 m of occupied stream and only 2 canals in 3262 m of unoccupied stream. Previous studies on beaver dispersion have also documented a preference for recolonizing areas where beavers have previously established (and abandoned) colonies^23, 24^.

Our observations showed that new canals were initiated during our field season at locations of particularly large and active trail systems (Fig. 4C). Google Earth imagery at other sites likewise provides other examples of trail systems becoming sites of initiation for canals (Extended Data Fig. 5B). These observations suggest that canals may serve as a more permanent record of successful colonization by previous colonies (Fig. 4D). While trails must be constructed from scratch each season when grassy vegetation re-grows, canal excavation provides long-term evidence and reinforcement of productive areas that are preserved beyond the typical lifespan of individual beavers (∼10–15 years).

## Discussion and Conclusions

By focusing on other relatively neglected aspects of beaver construction outside of dam building, particularly trails and canals, this novel dataset makes it possible to ask fundamental questions about *when* and *where* beavers decide to build. We observe trail clearing as the first construction and largest spatial-scale activity. Through active trail clearing, which precedes dam building (Fig. 3A), beavers can search for, access, and transport the vegetation needed for the construction of dams, lodges, scent mounds, and food caches. Though consistently occurring later, dam initiation did not follow at a consistent time after trail initiation. This variation likely reflects the natural variation in colony sizes and the proximity of building materials to the colony sites, where some sites required more trail construction or exploration time than others to source appropriate building materials.

Flow conditions in the stream influence *when* beavers build, beginning the trail-clearing necessary to establish a colony site. Beaver construction behavior has long been linked to cues associated with stream flow^15^. However, the results of this study emphasize the direct influence of flow on trail construction rather than dam construction. Moreover, the timing of the activity we observed is more selective than the indiscriminate “flow triggers building” association traditionally suggested. Waiting until the stream flow slows mitigates the likelihood of dam failure, as observed to occur during high-flow periods^25^—indeed, during our field season, a late-June storm event washed away incomplete dams at sites B and C that had been started that season, forcing those colonies to rebuild (Fig. 3D,G). The connection between flow and trail clearing suggests that beavers are not attempting and failing to build dams in unsuitable flow regimes. Instead, stream flow conditions provide them with information that lets them select conditions appropriate for building.

Preexisting canal architectures influence *where* beavers build. We found that beavers preferentially built trails close to canals. This proximity may help with the transportation of woody vegetation obtained by successful foraging along the trails.

Taken together, this evidence indicates that beavers respond to external environmental cues to inform both the spatial and temporal aspects of their construction behaviors. This study also emphasizes the importance of accessory architectures represented by trails and canals, which are the precursor structures required for the more commonly studied dams and lodges.

Trail building occurs in multiple phases (Fig. 3E). Early in the season, trail clearing can be associated with dam-building activities (Fig. 3A,D). Mid-season, total trail length plateaus or decreases as beaver trail-clearing activity decreases and vegetation overgrows previous trails. Late in the season, after the vegetation has reached its peak (Fig. 3F), trail clearing increases again, likely to support food caching necessary for winter survival. Vegetation quality may act as a late-season cue to prepare for winter.

A well-established trail system is likely to be repeatedly utilized by beavers if it provides a consistent source of food or construction materials. Conversely, unproductive trails become neglected and overgrown. Our observations suggest that over time, highly used trails can develop into canals (Fig. 4D, Extended Data Fig. 5B). Canals in turn may provide a key mechanism in the niche construction^9^ that promotes colony success over multi-year and multi-generational periods. Successful canals may undergo continual reinforcement and expansion from both beaver activity and water flowing through them; conversely, unsuccessful canals may silt up and dry, building up with nutrient-rich soil ideal for vegetation regrowth. We anticipate that a natural cycle of canal expansion and contraction (Extended Data Fig. 5E) could facilitate vegetation regrowth in these areas previously affected by canal construction, aided by an expanded riparian zone. In addition to facilitating access to terrestrial and aquatic resources, canals may result in a more hydrologically stable environment for the beaver colonies (Extended Data Fig. 8C, D). Coupled with dams, canals may help create a low-velocity environment, suited to beaver life-cycle requirements, by widening the riparian zone, deepening the stream depth, and bolstering the growth of riparian and wetland vegetation. Functionally, canals redirect water away from lodges, dams, and food caches during high-flow events, preventing large failure events of dams and increasing hydrodynamic stability of the damming complex. Canals further increase resilience to low-water events such as fires and droughts^26^. These factors imply the potential utility of canals as an indicator of landscapes conducive to long-term beaver occupancy and survival.

These observations suggest that architectural features may serve as a multigenerational coordination channel among beaver populations, indicating the potential suitability of a site to support beaver colonies and influencing their choice of where to establish new colonies. Recently, beaver dam analogs (BDAs) have been employed to mimic and attract beavers to new locations^27^ to mitigate the impacts of climate change across North America^28^. We suggest that the presence of canals, not just dams, would serve as a more effective way to engage with beavers and tap into their mechanisms of coordination for building. Influencing beavers on where to colonize will mitigate their conflict with humans while also enabling the many ecological benefits they provide.

## Supporting information

ExtendedData

## Methods

Using a combination of UAV photogrammetry, GIS (Geographic Information Systems), and in-field flow measurements, we reconstructed high-resolution 2D (∼1.5 cm/pixel) and 2.5D (∼4.5 cm vertical resolution) maps of beaver-modified habitats throughout the summer months. This section details site selection criterion; field data acquisition, including flight planning, ground control, and in-field flow measurements; orthomosaic generation in AgiSoft Photoscan; data extraction in ArcMap 10.7; and statistical analysis procedures.

### Site Selection

We selected three field sites with beaver-occupied lodges and a known history of dam building (Fig. 1C). The existence of lodges and the historical precedent of the dam building were determined from conversations with landowners and confirmed with an onsite visit to each site before data collection. All sites are located within the Blackfeet Indian Reservation (BIR) boundary. The BIR is located in the foothills of the Rocky Mountains in northwest Montana along the Canadian border (l48.746342°, -112.869043°). While beaver dams can be maintained for relatively long periods of time^29^, this work looks at colonies living under suboptimal conditions where the colony frequently builds new dams and rebuilds old dams. All sites chosen for this study were incised streambeds. The low-lying woody vegetation in the foothills of the Rocky Mountains allows for a relatively unobstructed view of the waterways while providing enough vegetation for beavers to build and sustain colony populations. First-order streams typically have one inlet and one outlet, simplifying the understanding of potential hydrodynamic cues that beavers may respond to.

Detailed descriptions of the field sites can be found in the supplementary materials.

**Ethical note:** Before data collection, this study was reviewed and approved by the Blackfeet Nation Institutional Review Board.

### Field data acquisition

Data was collected at an approximate frequency of once per week. Supplementary Table 2 provides specific days for which data was collected at each field site. Missions were planned according to weather conditions. Days with high wind conditions, extreme heat, or heavy rain were avoided for data collection. Flow measurements were taken upstream and downstream of the field sites on the same day UAV data was collected (unless otherwise noted).

### UAV and Ground Control

Prior to aerial data collection, ground markers were placed at consistent distances along the river at the selected field sites, using a Trimble GPS unit for precise latitude and longitude. These markers, placed in early May and removed in August, proved insufficient, so Google Earth Pro images with placemarks of common features were used for orthomosaic alignment. Two UAV models, the DJI Phantom Pro 4 and DJI Phantom 4 Advanced, were used, both with 1-inch CMOS sensors and 24 mm lenses producing 5472 x 3078 pixel images, including geolocation and altitude. DroneDeploy ensured consistent flight paths with a Nadir flight plan, achieving 75% front overlap and 65% side overlap at 12–16 m altitude for a resolution of 1.3–1.7 cm/pixel. Flights were conducted in the early morning to avoid battery overheating, with the drone equipped with a Marco Polo GPS tracker for each mission.

**Ethical note:** All flights were conducted or supervised by a licensed UAV pilot. To comply with the airspace regulations as defined by the United States Federal Aviation Administration, all flights were conducted within line of sight of the research team, a vertical ceiling of 120 m was respected, and the drone was not flown over national parks, people, private residences, or airfields. All private and public landowners granted permission and were notified as appropriate.

### In-field flow measurements

To measure fluid flow, we used a Marsh McBirney Flowmate 2000 with the velocity-area method to calculate the volumetric flow rate upstream and downstream of the field sites. The electromagnetic sensor measured water velocity, and these measurements were collected on the same day as the drone flights. Early season measurements were limited due to site accessibility, with upstream measurements at Sites B and C and downstream measurements at Site A.

### Vegetation phenology

To estimate the photosynthetic productivity of vegetation during our study, we calculated the following indices utilizing the reflectance of the visible green (ρ_green_), visible red (ρ_red_), and visible blue (ρ_blue_) bands of the RGB UAV imagery: Green-Red Vegetation Index (GRVI), Visible Atmospherically Resistant Index (VARI), and Green Chromatic Coordinate (GCC) (Extended Data Fig. 7).

GRVI is defined as^18^:

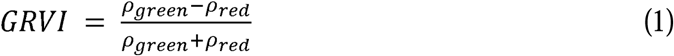

VARI, by subtracting out the reflectance of the visible blue (ρ_blue_), mitigates atmospheric effects^30^:

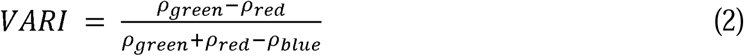

GCC is a measure of the relative amount of green vegetation in an image^31^:

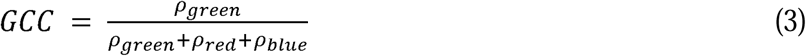

All three measures gave similar results (Extended Data Fig. 7); we show only GRVI in Fig. 3.

### Photogrammetry: Generating maps for UAV images

Following Agisoft Metashape Professional protocol, a batch process was built to repeat the steps of aligning drone images, optimizing the alignment, building a high-resolution dense cloud, creating a mesh for a 2.5D representation, generating texture, producing a tiled model, and generating a digital elevation model (DEM) and orthomosaic image, with each project saved and orthomosaics exported as TIFF files. This process took around 30 hours per scan, with each scan divided into chunks of approximately 200 images. Further details on model validation and feature extraction are in the supplementary materials.

### Analysis: ArcMap 10.7

The resulting orthomosaic was imported into ArcMap, an application of the ArcGIS Desktop package. To achieve alignment between subsequent orthomosaics, the GPS coordinates of the orthomosaics were removed and each orthomosaic was realigned and reshaped against satellite imagery using ground markers and stationary landmarks. Once each orthomosaic was aligned, dams and lodges resulting from beaver construction activities were then manually interpreted and represented as shape files of a characteristic shape, length, and spatial positioning. The orthomosaics and shape files are represented in the WGS 1984 ARC System Zone 12 coordinate system.

### Beaver damming complex features and classification

Shapefiles were created to classify different features of beaver damming complexes. A more detailed explanation of how shapefiles were used to define specific features can be found in the supplementary materials, along with a detailed annotation example (Extended Data Fig. 3A).

This work recorded surface-and-above-level changes made by beavers, focusing on above-ground features such as lodges, dams, canals, and trails, identified by experts in both fieldwork and remote sensing. However, the excavation depth of canals and ponds was not observed. We treat beaver damming complexes as quasi-2D and do not include measures of dam volume, pond depth, burrow volumes, or slopes in our analysis of these structures.

To estimate travel distances for beavers, we introduced a “distance along the stream,” reflecting their likely travel paths. The stream is defined by a manually drawn, geotagged line that helps in estimating distances within the riparian zone. The geotagged line is defined by uniformly spaced points 0.5 m apart. Each point is associated with a known latitude and longitude. Each beaver architectural feature—canals, trails, dams, and lodges—is associated with the nearest of these points, making it possible to calculate the distance between features along the river.

Width of the stream is defined for each point along the geotagged line based on where an orthogonal line at that point intersects each of the banks of the stream. Sinuosity^32^ is defined for each point along the geotagged line as (50 m)/*D*, where *D* is the Euclidean distance between the two points 25 m upstream and downstream along the geotagged line.

### Statistics

*Building initiation*: We performed multinomial exact tests (goodness-of-fit) to examine whether the first occurrence of construction of each type of structure for each colony is influenced by the date, GRVI, and/or flow rate of the river. We used our approximately weekly measurements of GRVI and flow rates at each site to linearly interpolate daily estimates for these variables. We then categorized these variables into bins and tested whether trail (or dam) initiation was equally likely to occur across bins. Though not strictly required for multinomial exact tests, we choose the breaks between bins such that the proportion of data points in each bin was similar, and made these choices without regard for the actual timing of construction initiation. We calculated the expected proportion of occurrences in a bin based on the relative proportion of time that the GRVI or flow rate of the river fell within the bin. For date, since beavers initiate construction early each summer, we only considered dates from May through mid-July (i.e. early season) as possible construction initiation dates.

We analyzed the distances between new dams and canals, and between trails and canals using a resampling approach. We compared the mean distance for each type of structure to the distribution of means derived from 1,000 simulated experiments in which the location of structures was randomized. We conducted these tests in two different ways: (1) In a pooled test, we considered all 19 new dams or 528 trails constructed by all colonies together, and the union of all territories (Fig. 4B–E). (2) In colony-by-colony tests, we considered only the dams or trails and the territory specific to each colony in turn (Extended Data Fig. 10A,C). In both cases, for each of the simulated experiments, the number of each type of structure was the same as in the real data, and locations were randomly chosen from within the corresponding territory. The resulting distributions of mean distances to a canal for each structure type and condition are therefore null distributions that would be expected if structures were located randomly. If the mean distance from dams or trails to canals was more extreme than extreme 5 percent of the associated null distribution (with two tails), we concluded that the structures were constructed non-randomly with respect to proximity to canals.

Stream characteristics at dam locations: We used similar resampling analyses to compare the mean stream {width, sinuosity} at dam locations with the null distribution of means expected if dam locations are chosen randomly. These tests were conducted under the pooled condition, considering all colonies and dams together.

### Study Limitations

This study tracks landscape changes associated with the results of beaver construction, not the act of construction. The (nocturnal) beaver was never directly observed in this study. This work does not estimate the number of individual beavers engaging in building behaviors nor how work is distributed amongst the colony members. The water depth of the streams was not collected and may be another potential factor influencing where beavers decide to build dams.

## Data and materials availability

The datasets generated during and analyzed during the this study are available in the Figshare repository, https://doi.org/10.6084/m9.figshare.26865394. This repository contains relevant data used in the analysis of this work. UAV-generated mosaics and shapefiles generated during and analyzed during this study are available from the corresponding author on reasonable request.

## Acknowledgments

We wish to express our deep gratitude to the Blackfeet Nation and the Blackfeet Nation Institutional Review Board for providing permission for this study to be conducted within tribal borders. We thank the private ranchers and farmers who allowed us access to their properties and wetlands; in particular the Michaels family, the Pilling family, the Kennedy family, the Magee family, the Skirka family, the Geer family, Mad Wolf Campground, and the Tatsey family. We would like to thank Blackfeet Fish and Wildlife for suggesting potential field site locations. We would also like to thank the many beaver practitioners, beaver believers, and beaver trappers who have offered their insights into beaver behavior. Furthermore, we are thankful to the Blackfeet community for encouraging this study to be conducted on a culturally significant species, in particular Dr. Brad Hall and Latrice Tatsey. We would also like to acknowledge Blackfeet Knowledge Keepers on sharing their insights working with Beaver, in particular, Helen Augare Carlson, Alicia YellowOwl, and John Murray. We would also like to thank Jeffrey Blossom from the Center for Geographic Analysis for providing access to and training on the necessary ArcGIS toolkits to conduct this study; Prof. Wade Campbell for sharing his expertise on drone mapping techniques and photogrammetry methodologies; and Dr. Shelly Lowe for her stable and persistent support of this research.

## Funding

Ford Foundation Predoctoral Fellowship

The National GEM Consortium Ph.D. Engineering and Science Fellowship

1665 Caleb Cheeshahteaumuch Predoctoral Fellowship

Professor Radhika Nagpal

## Author contributions

Conceptualization: JK

Methodology: JK, JW, AJ, DSH

Investigation: JK, AJ, DSH

Analysis: JK, HM, JW

Visualization: JK, HM, JW

Data Curation: JK, CC

Funding acquisition: JK

Supervision: JW

Writing – original draft: JK, HM, JW

Writing – review & editing: JK, HM, EF, JW

## Competing interests

Authors declare that they have no competing interests. Supplementary Information is available for this paper.

Correspondence and requests for materials should be addressed to.

Reprints and permissions information is available at www.nature.com/reprints.

## Notes

https://doi.org/10.6084/m9.figshare.26865394

### Competing Interest Statement

The authors have declared no competing interest.

