## ExtendedData for "Environmental cues influence timing and location of construction activity in a beaver damming complex"

### Extended Data

Jordan Kennedy[1]*, Helen McCreery[2], Audrey Jones[3], Dannette Spotted Horse[4], Candice Chen[5], Emily Fairfax[6], and Justin Werfel[1]

[1]Harvard University, Cambridge, MA 02138, USA

[2]Tufts University, Medford, MA 02155, USA

[3]Montana State University, Bozeman, MT 59717, USA

[4]Blackfeet Community College, Browning, MT 59417, USA

[5]Harvard College, Cambridge, MA 02138, USA

[6]University of Minnesota, Minneapolis, MN 55410, USA

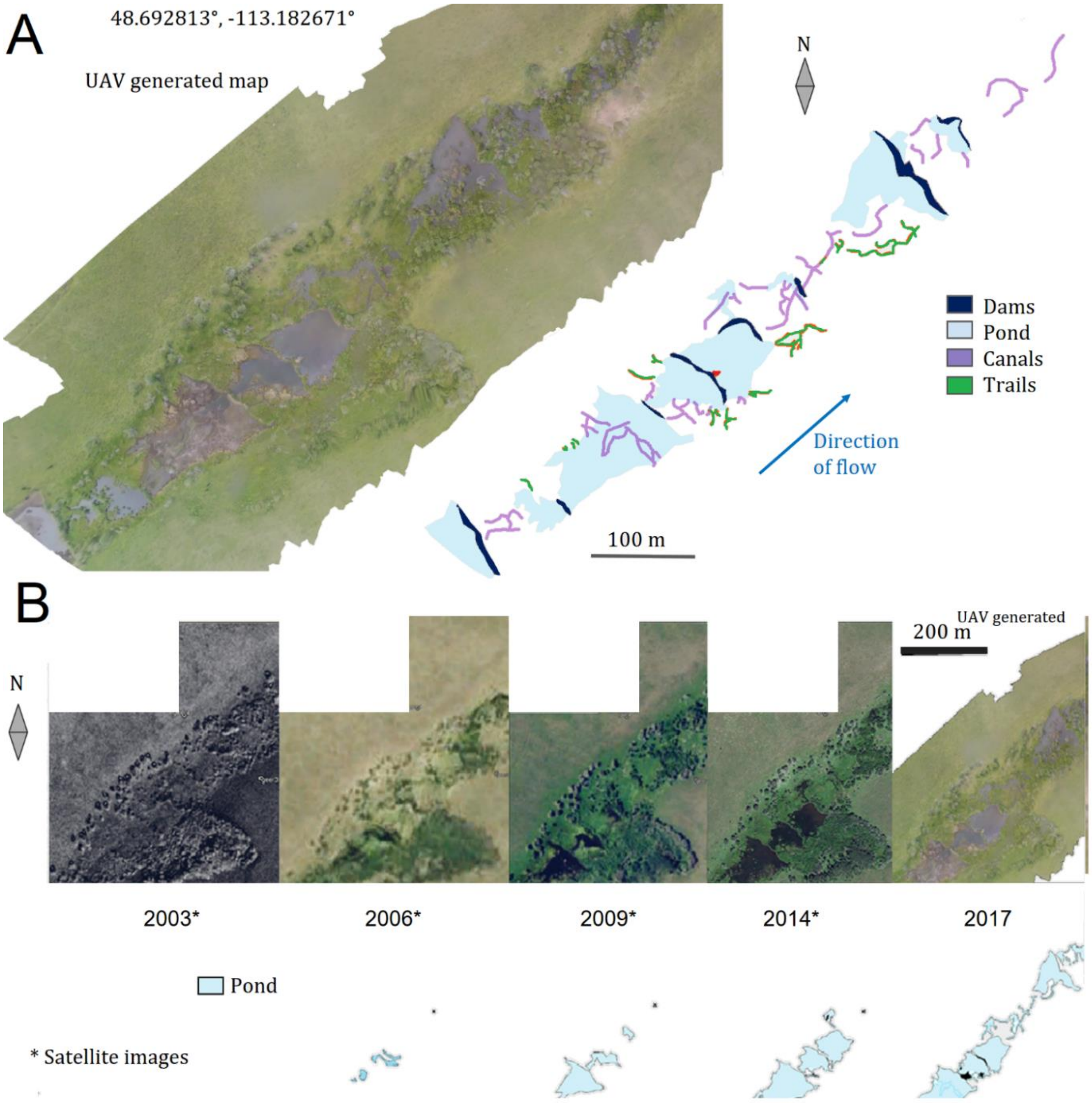


Extended Data Figure 1. **Spatial scale and temporal development of a damming complex.** (A) A 2017 UAV scan of a beaver damming complex on a private ranch within tribal boundaries. The scale demonstrates that a damming complex is orders of magnitude larger than any individual beaver, indicating the logistical engineering challenges beavers must overcome in building the complex. (B) Highlighting the exposed surface water over time emphasizes that a damming complex emerges over considerable temporal scales, frequently taking longer to build than a typical beaver lifetime, highlighting the organizational challenge of coordinating builders over large distances and multiple generations that beaver colonies must contend with. Images are from Google Earth[2] together with the UAV scan from (A).

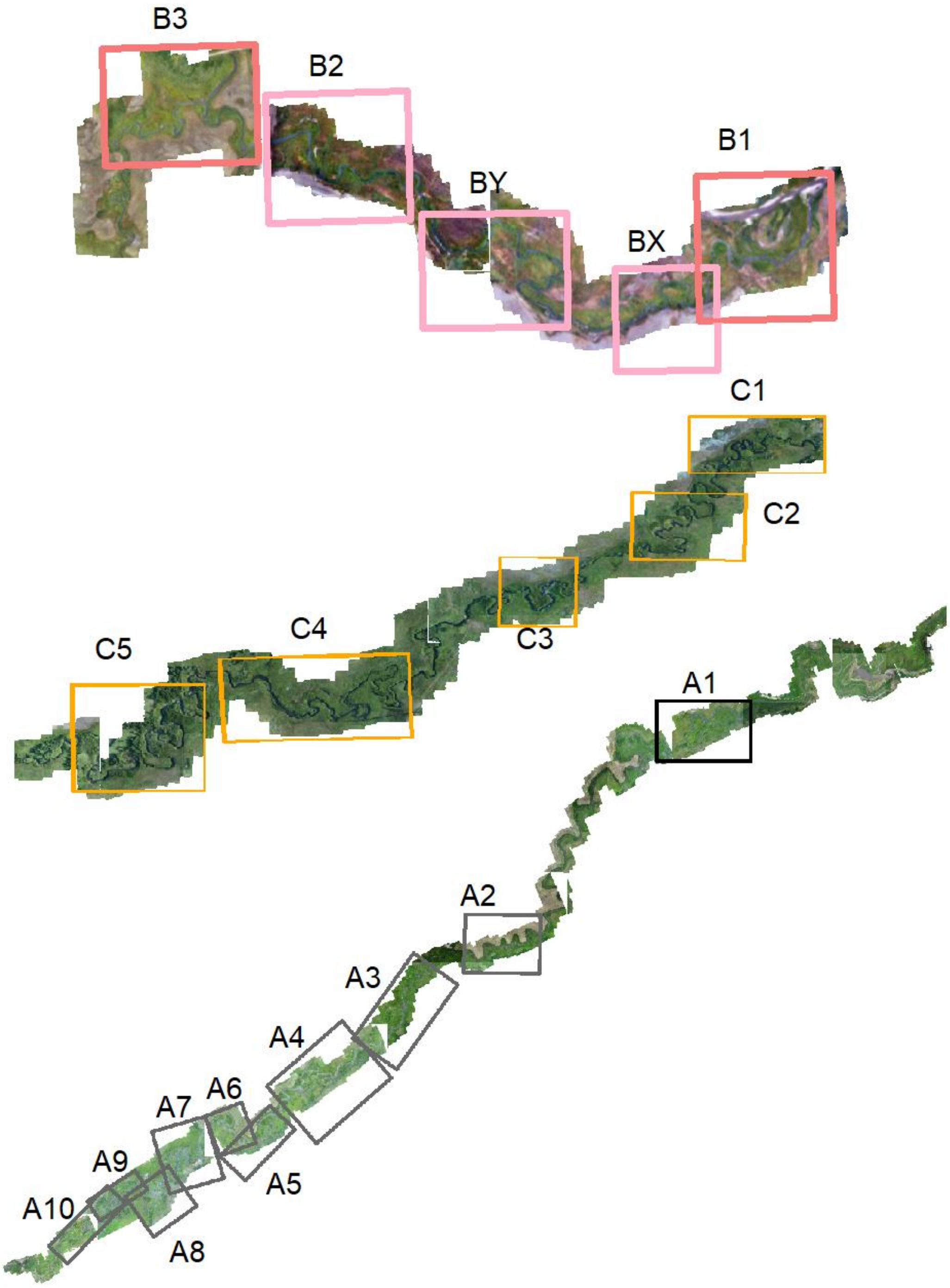


Extended Data Figure 2. **Colony boundaries.** Boundaries of each colony identified at each of the three field sites.

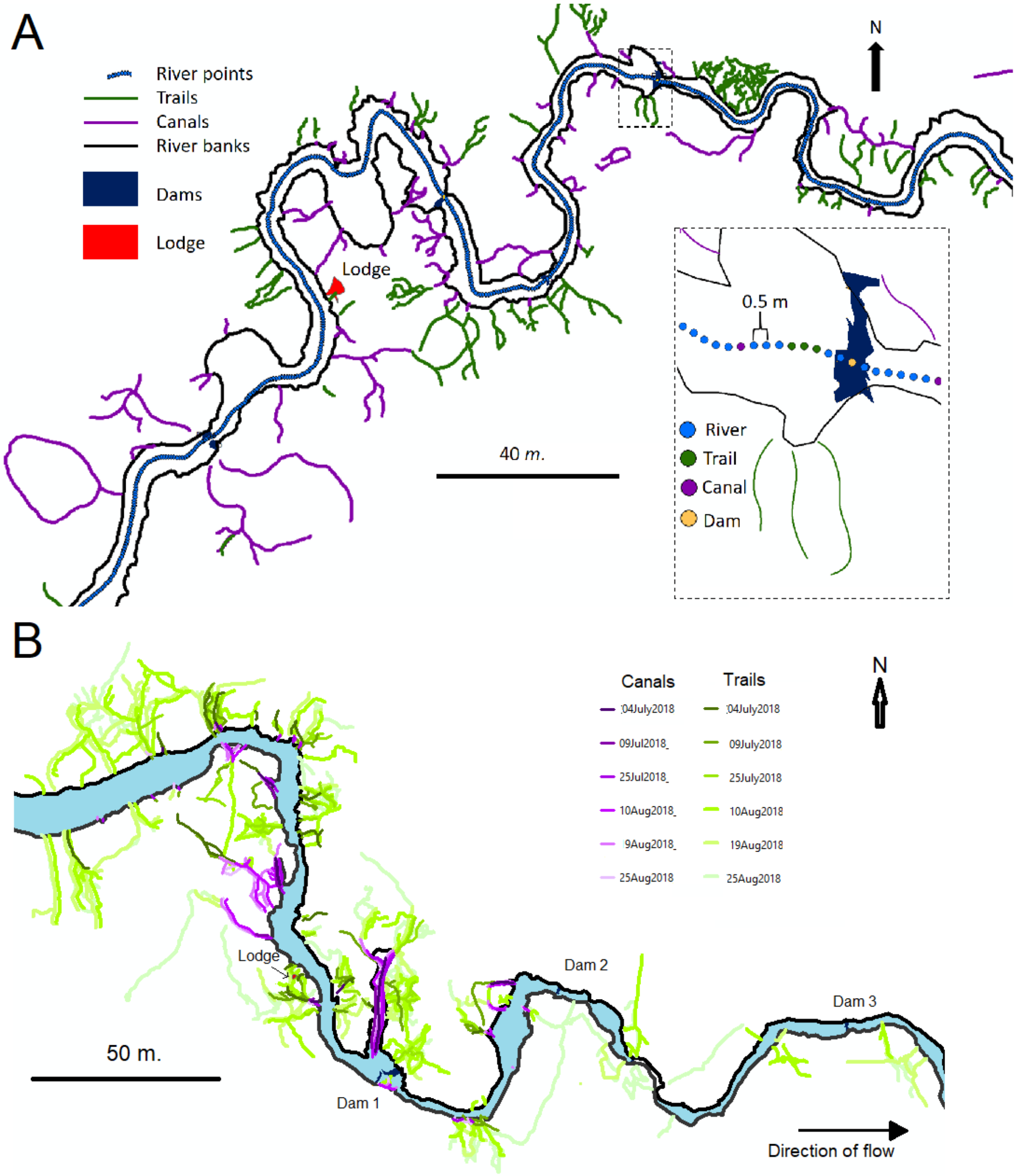


Extended Data Figure 3.**Annotated subregion of beaver colony spatially and temporally**. (**A**) Spatially, the river is represented by single points spaced 0.5 m. apart. Each river point has an associated latitude and longitude. Dams, lodges, and the points where canals and trails meet the stream are associated with the nearest river point. These associations are used to estimate the distance a beaver would travel along the river between two features. Trails, canals, and riverbanks are annotated with lines. These lines are characterized by shape, latitude, and longitude. Dams and lodges are characterized by area, perimeter, latitude, and longitude.(**B**) Temporally, colony B2 established a colony site later in the season. The largest dam (orange) is 1 m. from the nearest canal while the lodge (red) was established 34 *m*. from the nearest canal (purple). This colony was also not present during the 2019 UAV scan.

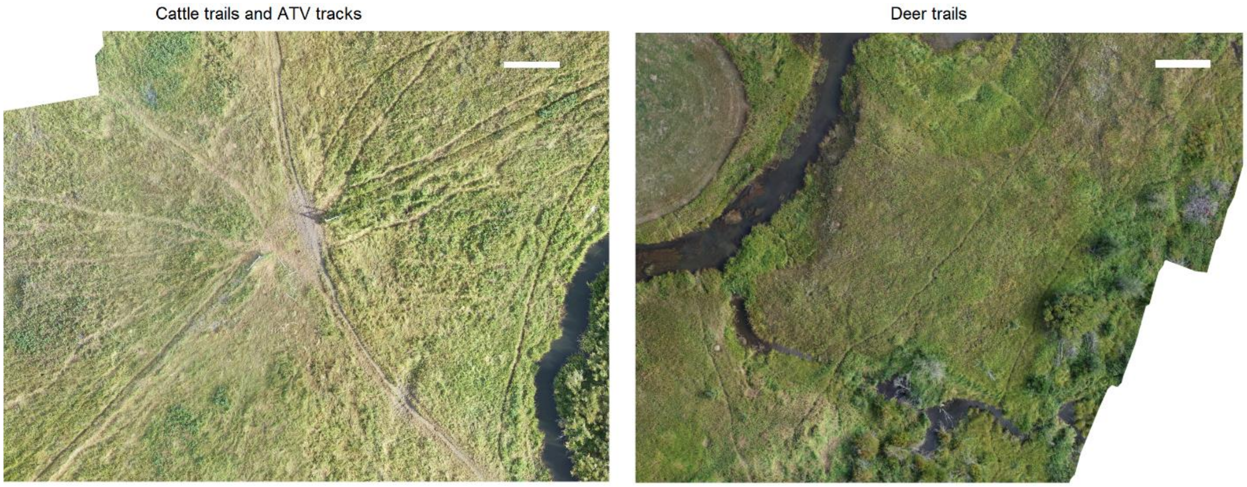


Extended Data Figure 4. **Landscape trails.** UAV images of other landscape changes associated with other species. (A) Cattle trails and ATV trails. (B) Deer trails. Scale bar: 5 *m*.

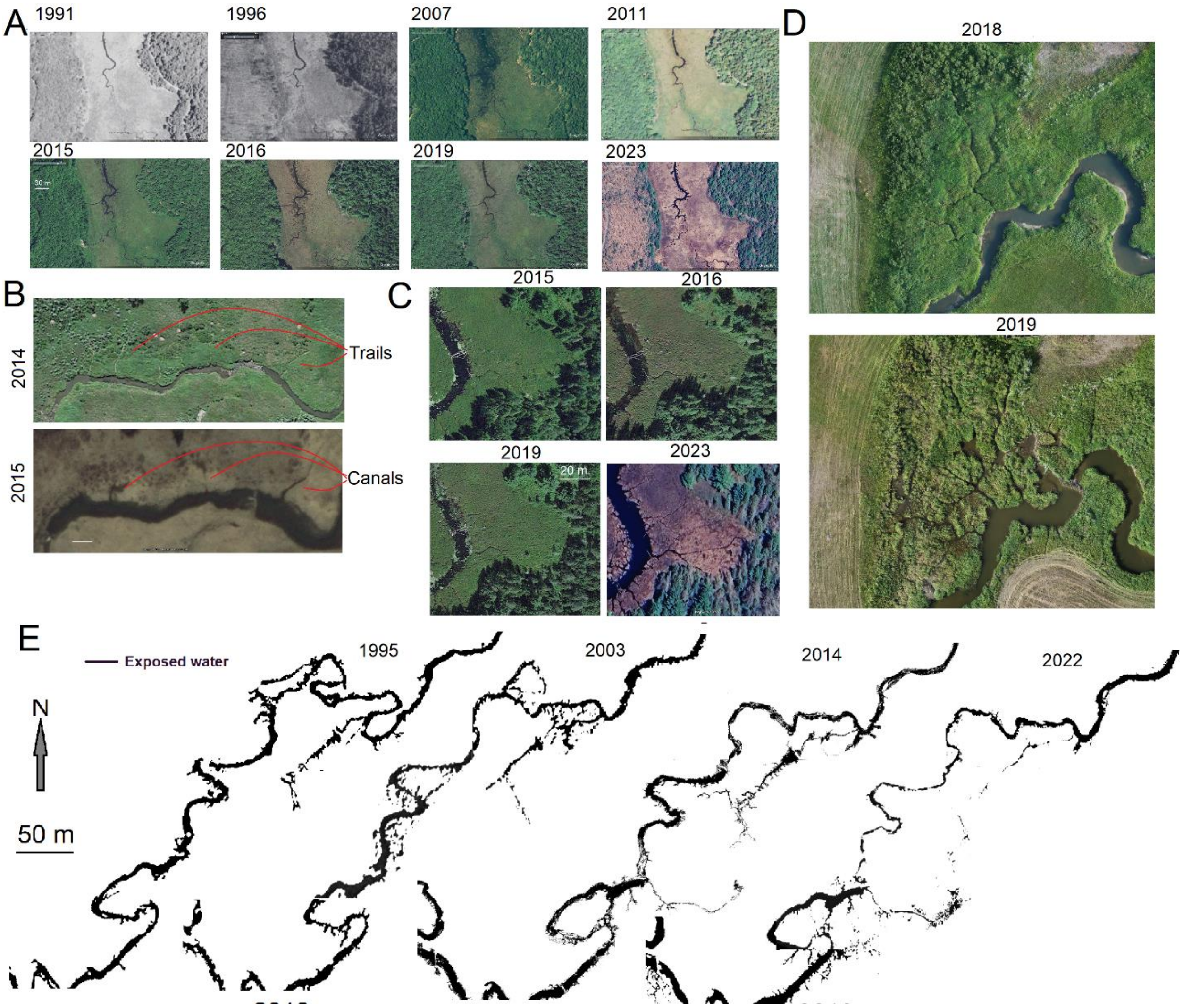


Extended Data Figure 5. **Trail to canal evolution.** (**A**) Example of beaver canal evolution from 1990 to 2022 at a location in Acadia National Park [14]. **(B)** Examples of canal initiation at locations of particularly active trail systems from 2014 and 2015 on the Blackfeet Indian Reservation in Montana [15]. **(C)** Canal evolution from 2015 to 2023 at another location in Acadia National Park [16]. All images are from Google Earth. (**D**) UAV drone imagery taken of a beaver damming complex with evidence of previous occupation (canals) that was unoccupied in 2018 and subsequently reoccupied in 2019. (**E**) Development of a canal system around a segment of river within Site A over a period of decades, showing notable expansion and contraction in different locations. Images are obtained from Google Earth and processed so that exposed water is shown in black[17].The waterways that branch away from the primary stream are canals.

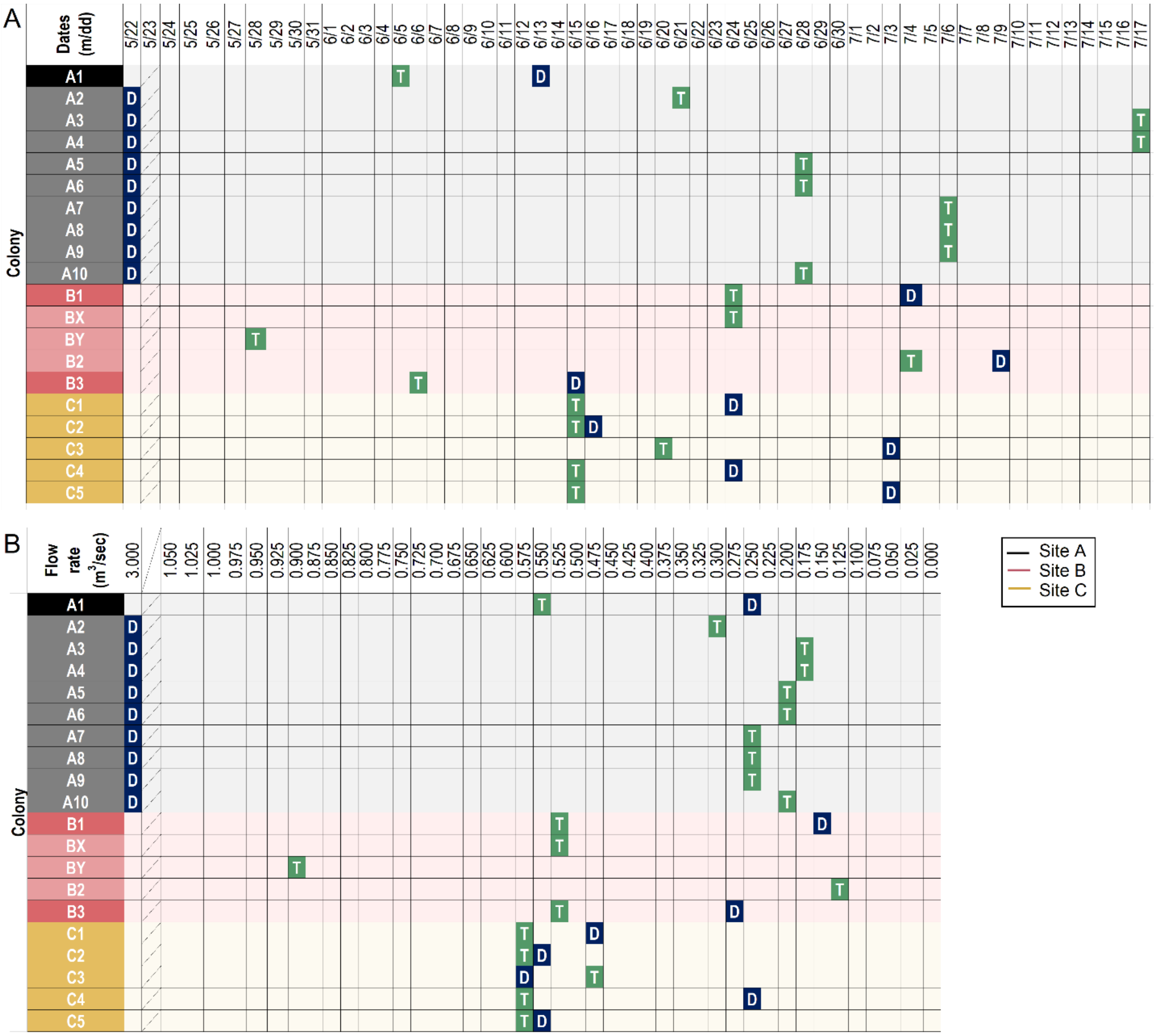


Extended Data Figure 6. **Initial observation of trails and dams for all 20 colonies.** For all 20 beaver complexes identified in this work, the (**A**) date and (**B**) flow rate when a trail system was first observed is marked with a “T”; dams are similarly marked with “D”. Colonies A2–A10 all had dams surviving from previous years. Colonies with no surviving dams needed to forage for material earlier in the season. Colony BX and BY abandoned their sites early in the season, building no dams.

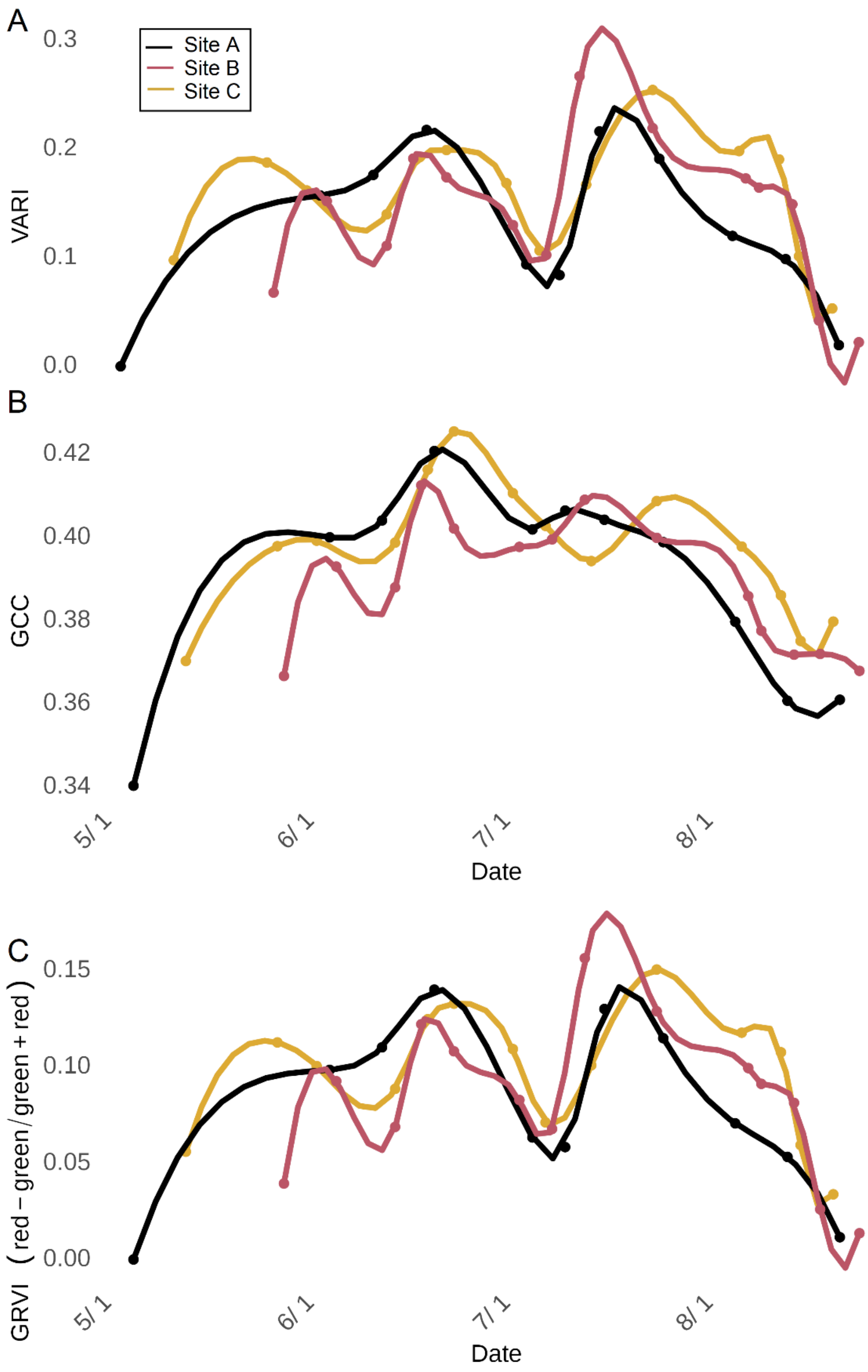


Extended Data Figure 7. **The phenological indices.** (**A**) VARI, (**B**) GCC, and (**C**) GRVI of all field sites.

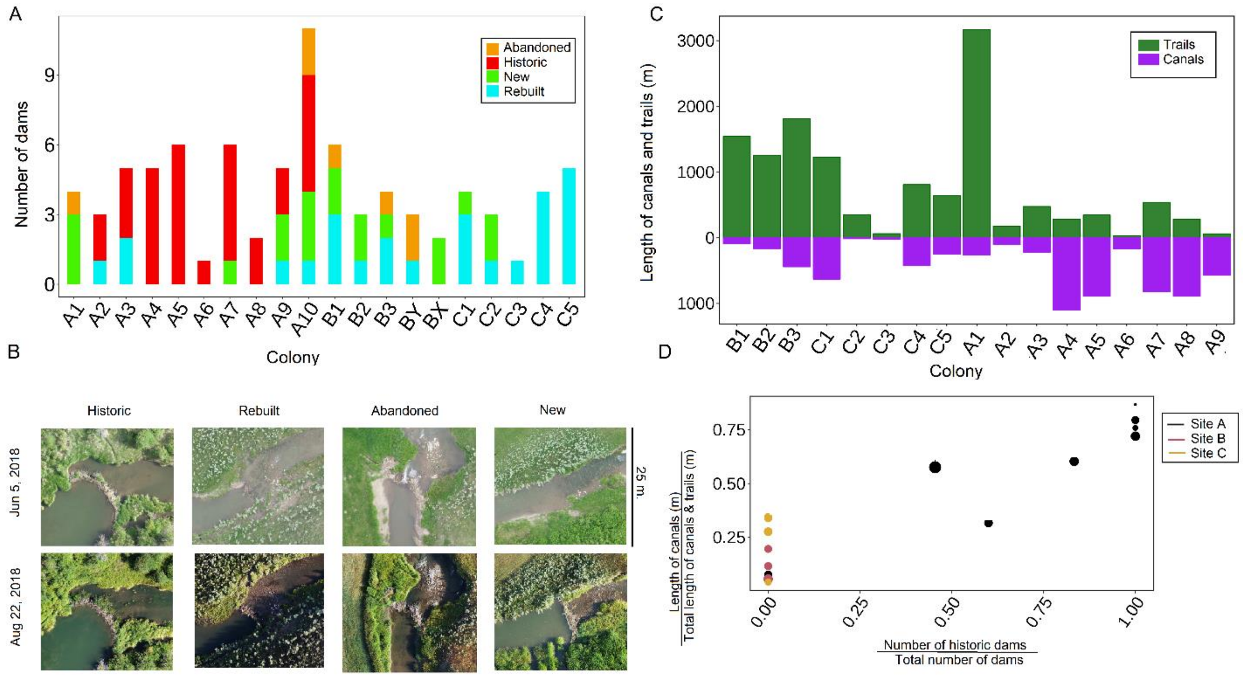


Extended Data Figure 8. **Architecture type and frequency.** (**A**) The number of dams of each category observed for each colony. In addition to those four categories, a single dam was observed to be built on a natural logjam; this dam was excluded from analyses. (**B**) The categories of dams in this dataset are historic, rebuilt, abandoned, and new. (**C**) Total lengths of all trails (green) and canals (purple) for the last scan for each colony. (**D**) Relationship between the ratio of the length of canals to the length of the trail-canal network and the number of historic dams to the total number of dams in each colony. The size of data points corresponds to the number of dams in each colony. Colonies with a greater fraction of canals are typically those with dams that survive the spring washout

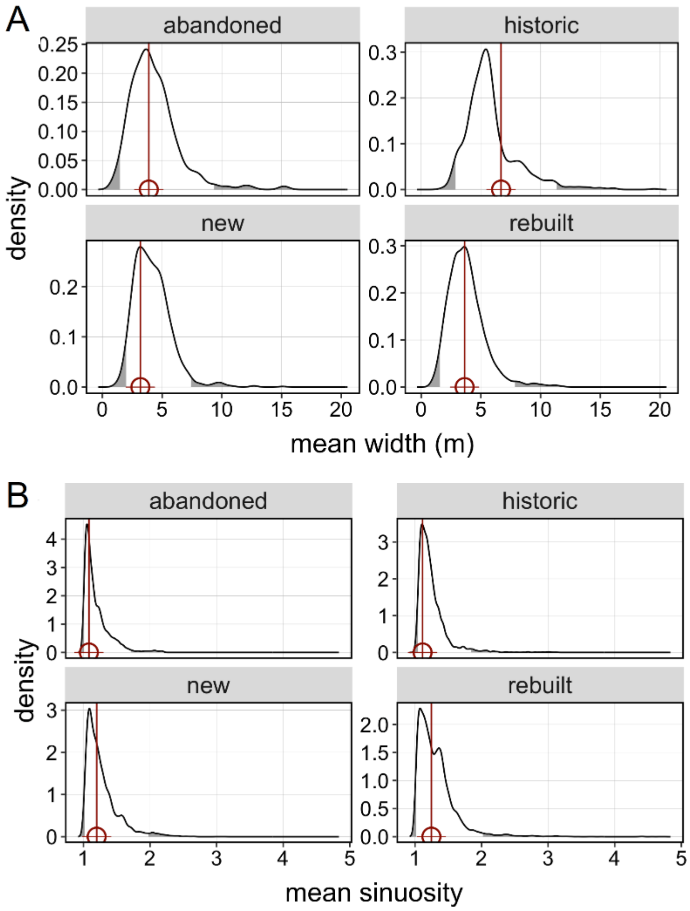


Extended Data Figure 9. **Resampling analysis broken down by dam type.** Resampling analysis for mean stream (A) width and (B) sinuosity at dam locations, for all four main dam types (all colonies pooled).

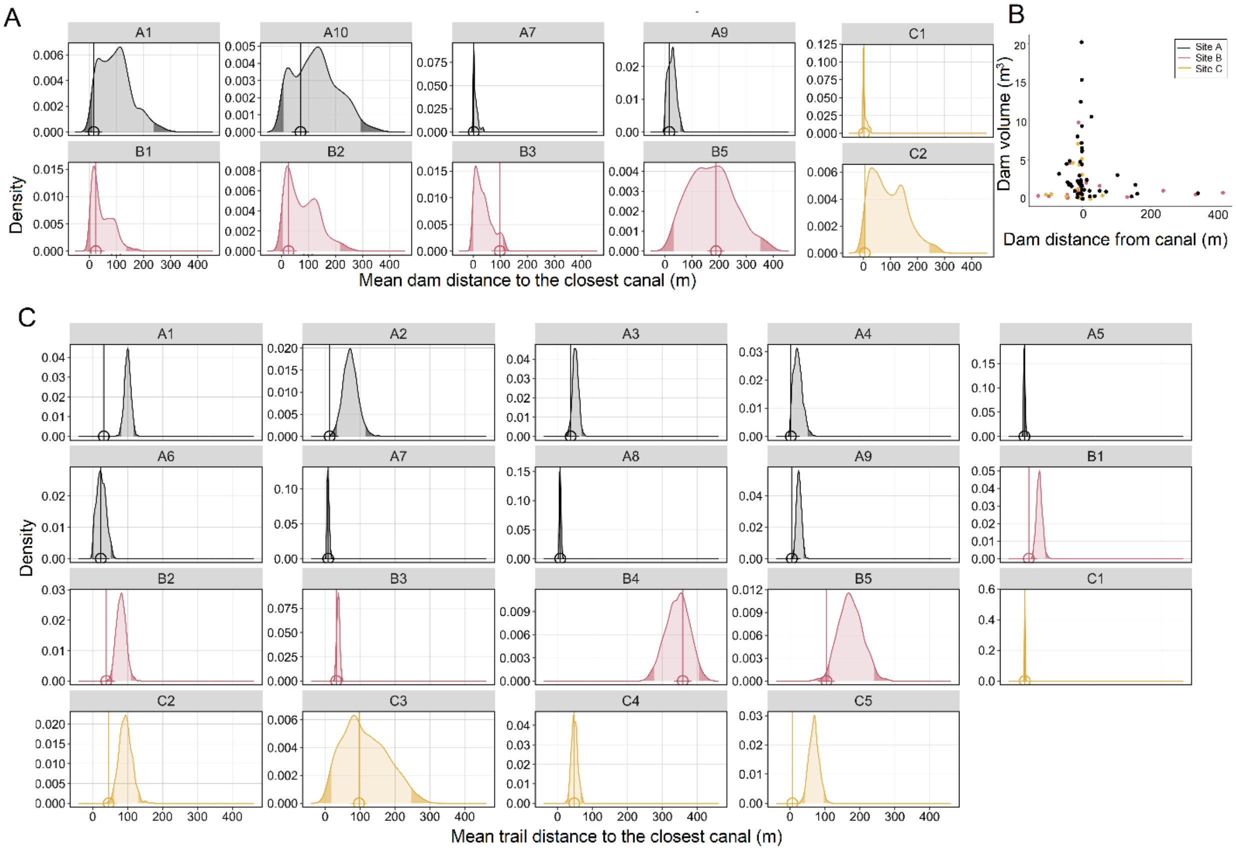


Extended Data Figure 10. **Proximity of beaver-built features to canals.** (**A**) Colony-by-colony resampling analysis for the mean distance between new dams and the nearest canal. (**B**) Dam volume plotted against location relative to the nearest canal (positive values correspond to downstream, negative values to upstream), for all dams. (**C**) Colony-by-colony resampling analysis for mean distance between trails and the nearest canal. The color of each point indicates the field site.
