## Supplement for "Environmental cues influence timing and location of construction activity in a beaver damming complex"

### Supplementary Information

Jordan Kennedy[1]*, Helen McCreery[2], Audrey Jones[3], Dannette Spotted Horse[4], Candice Chen[5], Emily Fairfax[6], and Justin Werfel[1]

[1]Harvard University, Cambridge, MA 02138, USA

[2]Tufts University, Medford, MA 02155, USA

[3]Montana State University, Bozeman, MT 59717, USA

[4]Blackfeet Community College, Browning, MT 59417, USA

[5]Harvard College, Cambridge, MA 02138, USA

[6]University of Minnesota, Minneapolis, MN 55410, USA

**Supplementary Methods**

Beaver damming complex

A beaver damming complex is built on spatial scales many times larger than a single beaver and over temporal scales much longer than the lifespan of a single beaver (Extended Data Fig. 1). Many beavers build the damming complex over multiple (frequently unrelated) generations to create, expand, and maintain the many structures that comprise the complex. The profound impact that beavers have on the reshaping of North American waterways has earned them the moniker of “Ecosystem Engineer”[1].

Beaver damming complexes contain seven distinct types of architectural elements, detailed in Supplementary Table 1.

UAV Missions

Drones (UAVs) were flown at an approximately weekly frequency at all field sites. Supplementary Table 2 gives a calendar of flight missions. The flight schedule was based on weather conditions, where unfavorable weather would prevent a mission from being flown.

Field site descriptions

All field sites are ecologically rich, with a plethora of species of large land mammals, such as wolves, black bears, grizzly bears, elk, deer, mountain lions, and moose; many bird species, such as nesting pairs of sandhill cranes, bald eagles, and hawks; and many aquatic species, such as leopard frogs, muskrats, otters, and multiple fish species. Low-lying woody vegetation primarily consisted of willow.

• Field site A (48.935030°, -113.116602°, first-order stream, 1315.212 *m.* elevation) covers a total area of 80 *ha.* along 10263 *m.* of stream. Site A is an active cattle ranch that is privately owned and sustainably managed. The land is mixed-use for both grazing and farming. This site rests upon the North Fork of the Milk River in the Milk River basin drainage, flowing primarily from west to east.

• Field site B (48.655272°, -112.750764°, first-order stream, 1204 *m.* elevation) covers a total area of 29 *ha.* along 3092 *m.* length of stream. Site B is a mixed-use field site owned both by the Blackfeet tribe and private ranchers who also farm the site for hay. Willow Creek, an S1 stream, flows primarily west to east through the Cut Bank basin drainage. While the area is largely grassland, woody vegetation is present, primarily consisting of Asher willow.

• Field site C (48.561810°, -113.058814°, first-order stream, 1520 *m.* elevation) covers a total area of 32 *ha.* on a total stream length of 4620 *m.* Like Site B, Willow Creek flows west to east in the Cut Bank basin drainage. Also like Site B, the Site C field site has multiple stakeholders, including tribal leases, private homeland plots, and active ranch and farmland.

Description of beaver colonies

Twenty beaver colonies were identified during this study: five at Site B, five at Site C, and ten at Site A (Extended Data Fig. 2; Supplementary Table 3). Site B and Site C consist entirely of new/emerging beaver colonies. At Site B, two colonies abandoned their territories early in the season, and a third colony migrated into the area mid-season. Of the ten colonies identified at Site A, nine had historic dams surviving from previous seasons.

*Determining beaver colony territory*

Extended Data Fig. 2 shows the upstream and downstream boundaries of each colony. These boundaries were defined by the locations of the most upstream and most downstream architecture (dam, trail, or scent mound, excluding the multi-year canals) for each lodge. Sites B and C had enough spacing between colonies that it was relatively straightforward to determine which colonies were responsible for building which structures. Site A had a clustering of colonies (A2–A10) close enough together that it was sometimes difficult to interpret which lodge a dam, canal, or trail belonged to. For this "supercolony" set of sites, each structure was assigned to the nearest lodge.

*2D Feature Extraction*

The orthomosaics were analyzed with ArcMap 10.7, an application of ArcGIS Desktop (Esri Company, Redlands, CA, USA). The results were exported and displayed in a WGS84 ARC System Zone 12 projection. Orthomosaics were manually aligned using ground control points and common geographical references.

Due to visibility conditions, variations in internal GPS readings of the UAV, variations in elevation take-off position of the drone, manual changes in flight pattern by the research team to account for a larger land area than anticipated, and wildlife interference (starlings flocking around the drone, hawks-dive bombing the drone, etc., resulting in a change in flight path), the final orthomosaics had distortions ranging from 1 cm to 1 m between flown missions.

Furthermore, in any encounter with large wildlife, such as bears or moose, the UAV was grounded until the animal left the area, resulting in slight variations in the final orthomosaic. To decrease the distortion of each orthomosaic generated by AgiSoft, each orthomosaic was subdivided into multiple smaller orthomosaics before being aligned in ArcMap. The GPS coordinates were removed from these smaller orthomosaics and manually redefined. All maps were aligned and scaled to their actual shape and size using known coordinates provided by the ground control points and preexisting satellite imagery.

Features were manually identified and annotated (Figs. 1, Extended Data Fig. 3).

*River annotation*

Riverbanks are annotated with line shape files. The center of a river is hand-drawn. The river is represented by individual points, spaced 0.5 *m*. apart. For each trail, dam, canal, and lodge, the closest river point is marked manually. The river points have an associated latitude and longitude, making it possible to calculate the distance along the stream between beaver-made features.

*Beaver dam features and annotation*

Dams are primarily woody structures originating from the edge of the stream and built inwards towards the center of the stream. Dams are represented as polygons. These polygons are quantified by their area and perimeter. Dams were further characterized by thickness and width. Thickness is defined as the length of a line segment passing through the center-most point of the dam, running parallel to stream flow, and reaching from one edge of the dam to the other. Width is defined as the length of the most upstream edge of the dam running from one bank to the other.

While a completed beaver dam is an obvious and abrupt feature on the landscape, the early stages of dam formation can present as ambiguous. To avoid ambiguity in dam identification, data analysis began with the final scan of each field site. A single shape was generated for each dam at each time point. This shapefile contains the area, length, and GPS coordinates of the dam. For consistency, dams were numerically identified with the most upstream dam at each site ordered as 1.

When identifying initial dam building, much of the initial dam construction activities occurred on dams that had been partially washed out and had remnants from the previous year. When new material added to these dams, that was considered the first initial dam observation in Fig 3A-C.  Area of the dams, prior to new builds, is captured in this work.

*Lodge features and annotation*

The majority of the lodges were identified at the beginning of the field season in May. These lodges were found by speaking with local landowners and tribal offices, and through scouting expeditions. River beavers, like the ones in this study, build their lodges on the riverbank. The lodge, like the dams, is comprised primarily of mud, rocks, weeds, and sticks. Some of these lodges are more like burrows than traditional lodges. These burrow lodges were identified later in the season and are characterized by underwater trenches perpendicular to the bank. Lodges were also represented in this study with shapefiles to characterize their size and location in absolute space and in relation to beaver dams.

Beaver lodges are also represented as polygon shape files. However, as all field sites in this study were on rivers, most of the beaver lodges were also part of burrows. Therefore, the size of the lodge that could be seen from the air did not give a good estimate for the total lodge size as much of the lodge was underground. Burrow entrances were identified by entrance canals which were visible through the water on the bottom of the stream.

*Trail identification and annotation*

Beaver trails require a relatively high resolution to be seen in satellite or UAV imagery. The UAV imagery in this study provides a fine enough resolution to resolve trails, while the majority of satellite imagery accessed in this study did not (with some exceptions, e.g.,Extended Data Fig. 5B).

As a wild space, many species, including beavers, utilized the habitat. Many of these species, including cattle, horses, elk, deer, and ants, also impacted the landscape through trail-making (Extended Data Fig. 4). Therefore beaver trails needed to be distinguished from those made by other animals for this study.

Beaver trails are distinct from other trails in that they always start at the stream bank and initially run perpendicular to the stream (cf. Figs. 1*A.v.*, 2, 4). These trails also branch, which is not observed in trails made by the other species noted above. Additionally, beaver trails are only observed near other beaver-made architectures (dams, lodges, and canals). Beaver trails are also cleared of vegetation, making them distinct from other trail systems, which are typically characterized by a flattening of vegetation but not removal of vegetation. Lines are used to represent trails, producing a characteristic length. Trail width variation is not reported in this work. Future analysis may include trail width as a proxy for frequency of use, where more used branches of a trail indicate heavier traffic.

*Canal features and annotation*

Canals are beaver-excavated channels extending from the edge of the stream. Canals are defined as ditches filled with water. Empty ditches, while abundant in this dataset, particularly early in the season, were not annotated as canals. Empty ditches are distinct from overland trails in that, unless they are very old, they do not exhibit the branching characteristic of trail systems. Additionally, they are frequently rich in vegetation. Late-stage canals, which run perpendicular to the stream path, are visually easy to detect, being filled with water and extending several meters away from the edge of the stream. Canal width, which can vary as it becomes filled with water, was not captured in this work.

Early-stage canals, often present as minor indentations in the stream bank, can develop from beaver slides, places where beavers frequently enter and exit the water.
Active canals, like trails, are represented with lines.

### Supplementary Tables

Supplementary Table 1. **A summary of form and function of beaver-built architectures.**

| **Beaver-built architectures** | | |
|---|---|---|
| **Category** | **Structure** | **Description** |
| **Seasonal-Lifetime** | Food cache | A food cache is a collection of woody vegetation to be eaten during the winter months. As beavers do not hibernate, they need a food source to maintain the colony throughout the winter months. In colder regions, the food cache is weighed down by rocks to the deepest part of the pond. This enables the beaver to swim under the ice all winter long to retrieve food[3-5]. |
| | Lodge | All activities in a beaver colony are centered around the lodge. The lodge is where beavers eat, sleep, and raise their young. Beavers will often abandon their lodges and rebuild new ones nearby. It is not uncommon for a colony to rebuild a new lodge annually. The lodge, like the dam, is primarily made out of mud, woody vegetation, and rocks. The lodge can take three forms: the classic lodge in the middle of the pond, burrows in the sides of river banks, or hybrid structures combining features of both[6,7]. |
| | Scent mounds | Scent mounds are mounds of dirt constructed by beavers. As the name implies, scent mounds are scented from the castor sacs of the dominant adults. These scent mounds serve as territorial markers and are aggressively maintained and guarded by colony members from outside beavers[8,9]. |
| | Trails | Beaver-cleared trails arise as beavers transport material over land. As woody vegetation is cleared, a path is created so that beavers can continue to travel inland to retrieve materials. These trails have been reported to be dozens of meters in length[10]. |
| **Multi-generational** | Canals | In order to transport materials that would otherwise be too cumbersome to move over land, beavers excavate canals. In this way, beavers can extend their ponds and create "highways" for material transport by enabling them to float woody vegetation back to a lodge, food cache, or dam[11]. |
| | Dams | Perhaps the most iconic of beaver-built structures is the beaver-made dam. Beaver dams are porous structures built of locally available materials. Inspecting a dam will give the viewer a detailed sampling of local vegetation. The final structure spans streams, rivers, and canals. Beaver dams have been known to be built out of a diverse range of materials, including road signs, plastic containers, Styrofoam, animal remains[12], bags of money[13], and even beaver traps. |
| | Beaver pond | The pond behind a dam provides protection and creates an environment that supports beaver lifecycle activities. |

Supplementary Table 2. **Summary of flights at each of the field sites.**

| Site A | Site B | Site C |
|---|---|---|
| - | May 5th, 2018 | - |
| May 6th, 2018 | - | May 6th, 2018 |
| May 16th, 2018 | May 15th, 2018 | May 14th, 2018 |
| May 22nd, 2018 | May 20th, 2018 | May 21st, 2018 |
| - | May 29th, 2018 | May 28th, 2018 |
| June 5th, 2018 | June 6th, 2018 | June 3rd, 2018 |
| June 13th, 2018 | June 15th, 2018 | June 15th, 2018 |
| June 21st, 2018 | June 19th, 2018 | June 20th, 2018 |
| June 28th, 2018 | June 24th, 2018 | June 24th, 2018 |
| July 6th, 2018 | July 4th, 2018 | July 3rd, 2018 |
| - | July 9th, 2018 | July 8th, 2018 |
| July 17th, 2018 | July 14th, 2018 | July 14th, 2018 |
| July 26th, 2018 | July 25th, 2018 | July 25th, 2018 |
| - | Aug 8th, 2018 | Aug 7th, 2018 |
| Aug 6th, 2018 | Aug 10th, 2018 | Aug 13th, 2018 |
| Aug 14th, 2018 | Aug 15th, 2018 | Aug 15th, 2018 |
| Aug 22nd, 2018 | Aug 19th, 2018 | Aug 21st, 2018 |
| - | Aug 25th, 2018 | - |

Supplementary Table 3.**Table of identified beaver colonies during the 2018 field season.**

| Site | Colony | Complex type | Status | Established | Latitude | Longitude |
|---|---|---|---|---|---|---|
| Site A | A1 | New | Active | Prior | 48° 56' 38.2812" N | 113° 5' 56.4" W |
| Site A | A2 | Historic | Active | Prior | 48° 56' 8.6604" N | 113° 6' 40.5756" W |
| Site A | A3 | Historic | Active | Prior | 48° 56' 1.9824" N | 113°7' 4.4256" W |
| Site A | A4 | Historic | Active | Prior | 48° 55' 51.4632" N | 113° 7' 25.7196" W |
| Site A | A5 | Historic | Active | Prior | 48° 55' 42.6612" N | 113°7' 35.3532" W |
| Site A | A6 | Historic | Active | Prior | 48° 55' 44.6772" N | 113°7' 44.8068" W |
| Site A | A7 | Historic | Active | Prior | 48° 55' 39.702" N | 113° 7' 52.986" W |
| Site A | A8 | Historic | Active | Prior | 48° 55' 35.994" N | 113°7' 59.3688" W |
| Site A | A9 | Historic | Active | Prior | 48° 55' 35.652" N | 113°8' 4.9668" W |
| Site A | A10 | Historic | Active | Prior | 48° 55' 29.766" N | 113°8' 15.6012" W |
| Site B | B1 | New | Active | Prior | 48° 39'21.88800" N | 112°15'24.8400" W |
| Site B | B2 | New | Active | Late season | 48° 39'27.61200" N | 112°14'13.9200" W |
| Site B | B3 | New | Active | Prior | 48° 39'31.64400" N | 112°14'17.5200" W |
| Site B | Bx/B4 | New | Abandoned | Prior | 48° 39'16.56000" N | 112°15'03.6000" W |
| Site B | By/B5 | New | Abandoned | Prior | 48° 39'23.86800" N | 112°14'52.0800" W |
| Site C | C1 | New | Active | Prior | 48° 33' 50.7456" N | 113° 3' 17.3556" W |
| Site C | C2 | New | Active | Prior | 48° 33' 45.0216" N | 113° 3' 25.5276" W |
| Site C | C3 | New | Active | Prior | 48° 33' 39.6396" N | 113°3' 37.5228" W |
| Site C | C4 | New | Active | Prior | 48° 33' 30.2796" N | 113°3' 57.5028" W |
| Site C | C5 | New | Active | Prior | 48° 33' 26.0388" N | 113° 4' 15.7908" W |

**Supplementary Results**

Due to the near-extinction levels of beavers across North America, we have the opportunity to observe how landscapes change as beavers recolonize their previous habitats, from which they have been missing for decades to centuries. Extended Data Fig. 5 show that the occupation of a site is not a steady linear expansion. Rather, beaver-occupied habitat undergoes cycles of colonization and abandonment. In this cycle we can expect to see an expansion of the riparian zone when beavers are present and a contraction as beavers abandon the site. Site abandonment and the consequential recolonization on a multi-year cycle by beaver colonies is a well-observed phenomenon that is not well-studied. Site abandonment may potentially be attributed to factors including 1) vegetation changes, 2) hydrodynamic changes, 3) predator influences, 4) site contamination with beaver waste, and/or 5) beaver-to-beaver site competition.

Five dam types were identified in this study (Extended Data Fig. 8 A,B). Historic dams are defined as dams that were not washed out during the spring flood. Instead, these dams survived from previous years, and were actively maintained by beaver colonies throughout the field season. 2) Rebuilt dams are defined as dams that had previously been washed out during the spring flood and were rebuilt by beavers in the same location. These dams were visually identifiable as remnants of their structure could be seen along the stream bed. 3) New dams are defined as dams with no evidence of previous dams being built in that location. 4) Abandoned dams are defined as those that had previously been washed out during the spring flood but for which no rebuilding activity occurred. Like rebuilt dams, these were visually identifiable from their remnants. 5) In one case, a natural log jam in the river was later built on by beavers to be modified into a dam. The natural log jam had woody material aligned perpendicular to the river's flow; after modification into a beaver dam, material was oriented in many different directions. This unique dam was excluded from the analyses in the main paper.

By plotting the ratio of historic dams to the total number of dams in a colony against the ratio of the length of canals to the combined length of trails and canals, we can see that colonies with many historic dams also have a higher proportion of canals (Extended Data Fig. 8 C,D). New and emerging colonies have a lower ratio of canals to trails. This correlation is consistent with the idea that canals help stabilize a damming complex, making them more resistant to high-flow events.

*Resampling analysis*

Extended Data Fig. 9A shows the mean width of the river for dams of each of the four main (non-logjam) types, with dams and territories for all colonies pooled together. No dam type is built at an unusual width compared to the respective null distribution. However, the widths at locations of different dam types do differ (Kruskal-Wallis: $X^2 = 17.4$, df = 3, $p < 0.001$). Specifically, historic dams are at significantly wider locations than both new dams and rebuilt dams (Dunn post-hoc tests with Bonferroni correction: historic dams vs new dams, $p < 0.001$; historic vs rebuilt dams, $p < 0.01$; all other comparisons, $p > 0.05$). This result is consistent with the previously reported proposal that dams have the effect of widening the stream at their location over time[18].

Extended Data Fig. 9B shows the mean sinuosity of the river for dams of each of the four main types, with dams and territories for all colonies pooled together. No dam type is built at an unusual sinuosity compared to the respective null distribution. As described in the main text, when we perform pooled analyses considering all colonies together, we find that trails—but not dams—are constructed significantly closer to canals than expected by chance. To confirm that these results are present *within* colonies, and are not artifacts of inter-colony variation, we performed the analysis separately on each colony's territory.

Ten colonies built new dams during the field season (Extended Data Fig. 10A). Of these, one (C2) built dams in the left tail of the null distribution for distance from canals for that colony. In general, however, the observed mean distances from new dams to canals fell well within the null distributions. Based on this analysis, and consistent with the pooled analysis, there is no clear effect of distance to canals on locations of new dams.

All twenty colonies built trails during the field season (Extended Data Fig. 10C). Eight of these (A1, A2, A4, A9, B1, B2, C2, C5) built trails in the extreme lower tail of the null distribution for that colony, indicating that trails were constructed far closer to canals than expected by chance. This pattern also occurred at four more colonies (A3, B3, B5, C3), though to a less extreme extent. In the territories of four other colonies (A5, A7, A8, C1) there were so many canals that every point along the river was close to a canal, making the null distribution very narrow near 0, so that we were unable to test the question properly in these cases. One colony (A10) had extensively modified its territory over the course of decades so that the stream was diverted in part into many smaller side channels, with no trails at the end of the season visible along the main channel; thus A10 was excluded from this analysis. For three colonies (A6, B4, C4), the pattern observed in the pooled analysis was not clearly reproduced with our data. We therefore conclude, both for dams and for trails, that the result reported in the pooled analysis (Fig. 4Bi-iv) is not an artifact of differences among colonies.

We observe a pattern of larger dams being built closer to canals, with the largest being very close to a canal (Extended Data Fig. 10B). This observation is consistent with the idea that canals facilitate access to woody vegetation, such that bringing more vegetation to construct a larger dam is easier at locations closer to canals.
